# Robust high throughput prokaryote *de novo* assembly and improvement pipeline for Illumina data

**DOI:** 10.1101/052688

**Authors:** Andrew J. Page, Nishadi De Silva, Martin Hunt, Michael A. Quail, Julian Parkhill, Simon R. Harris, Thomas D. Otto, Jacqueline A. Keane

## Abstract

The rapidly reducing cost of bacterial genome sequencing has lead to its routine use in large scale microbial analysis. Though mapping approaches can be used to find differences relative to the reference, many bacteria are subject to constant evolutionary pressures resulting in events such as the loss and gain of mobile genetic elements, horizontal gene transfer through recombination and genomic rearrangements. *De novo* assembly is the reconstruction of the underlying genome sequence, an essential step to understanding bacterial genome diversity. Here we present a high throughput bacterial assembly and improvement pipeline that has been used to generate nearly 20,000 draft genome assemblies in public databases. We demonstrate its performance on a public data set of 9,404 genomes. We find all the genes used in MLST schema present in 99.6% of assembled genomes. When tested on low, neutral and high GC organisms, more than 94% of genes were present and completely intact. The pipeline has proven to be scalable and robust with a wide variety of datasets without requiring human intervention. All of the software is available on GitHub under the GNU GPL open source license.

**DATA SUMMARY:**

1. The assembly pipeline software is available from Github under the GNU GPL open source license; (url - https://github.com/sanger-pathogens/vr-codebase)
2. The assembly improvement software is available from Github under the GNU GPL open source license; (url - https://github.com/sanger-pathogens/assembly_improvement)
3. Accession numbers for 9,404 assemblies are provided in the supplementary material.
4. The *Bordetella pertussis* sample has sample accession ERS1058649, sequencing reads accession ERR1274624 and assembly accessions FJMX01000001-FJMX01000249.
5. The *Salmonella enterica subsp. enterica serovar* Pullorum sample has sample accession ERS1058652, sequencing reads accession ERR1274625 and assembly accession FJMV01000001-FJMV01000026.
6. The *Staphylococcus aureus* sample has sample accession ERS1058648, sequencing reads accession ERR1274626 and assembly accessions FJMW01000001-FJMW01000040.

**I/We confirm all supporting data, code and protocols have been provided within the article or through supplementary data files**.☑

**IMPACT STATEMENT:** The pipeline described in this paper has been used to assemble and annotate 30% of all bacterial genome assemblies in GenBank (18,080 out of 59,536, accessed 16/2/16). The automated generation of *de novo* assemblies is a critical step to explore bacterial genome diversity. MLST genes are found in 99.6% of cases, making it at least as good as existing typing methods. In the test genomes we present, more than 94% of genes are correctly assembled into intact reading frames.

## INTRODUCTION

***The rapid reduction in the cost of whole genome sequencing has made it feasible to sequence thousands of prokaryotic samples within a single study (Chewapreecha et al. 2014; Nasser et al. 2014;Wong et al. 2015). Many bacteria acquire genetic material through horizontal gene transfer when different strains recombine (Croucher et al. 2011). Mobile genetic elements such as phage, plasmids and transposons, by their very nature, are the most variable part of the genome, enabling rapid exchange of genetic material between isolates. They are known to carry antibiotic resistance and virulence genes, and so are some of the most biologically interesting parts of the genome (Medini et al. 2005). Identifying lost sequences and genes is also biologically important as this can signal host or environment adaptation (Klemm et al. 2016). Though reconstructing the sequence (de novo assembly) and performing annotation is a more complex process than performing a mapping based approach, it will: (1) generate sequence not in the reference genome (variable accessory genome (Page et al. 2015)), (2) resolve deletions which generate errors in mapping based approaches, (3) find signatures of recombination (Croucher et al. 2014), and (4) enable the community to work with a full sequence for bottom up analysis from public databases, rather than SNP lists***.

Although de novo assembly is computationally challenging (Pop 2009) it has many advantages over mapping based approaches. One of the fundamental limitations of de novo assembly is that any repetitive regions within the genome that exceed the length of the library fragment size prevent a complete de novo assembly from paired end reads. However, the most cost effective, and hence most common, sequencing method involves sequencing the ends of short DNA fragments (<1000bp). When a repeat region is larger than the fragment size, the assembler cannot unambiguously reconstruct the underlying sequence, so a break is introduced. This challenge has been addressed in a number of different ways. Automatically tuning parameters and configurations can produce improved assemblies, such as using RAMPART (Mapleson et al. 2015). The MetAMOS pipeline (Treangen et al. 2013) uses multiple different assemblers and picks the best result, however it takes over one month to assemble a single bacterial genome which makes it computationally unfeasible to run on a large number of samples. Assemblies may be improved using wet lab methods (Puranik et al. 2015), such as using capilary sequencing to extend over gaps, optical mapping, or additional long insert mate pair libraries, however these approaches are low throughput and prohibitively costly. Several software tools exist to perform scaffolding (Hunt et al. 2014), and automated gap closing (Tsai et al. 2010; Boetzer & Pirovano 2012) which improve the de novo assembly step. The annotation of bacterial genomes can be programatically performed using a number of automated tools (Seemann 2014; Mitchell et al. 2015). Although we have seen a commoditisation of sequencing technologies due to rapidly decreased costs, the generation of annotated genomes, and deposition of those to the public archives (EMBL/GenBank), can be a very time consuming and laborous process, so is rarely performed (Pirovano et al. 2015). Taking the Salmonella genus as an example: of the 44,920 WGS samples sequenced, only 4,451 (9.9%) have had assemblies depositied in GenBank (accessed 5 May 2016).

To overcome these challenges, we have created a reliable assembly and improvement pipeline that consistently produces annotated genomes on a large scale ready for uploading to EMBL/EBI. To date, 18,080 de novo assemblies have been created and submitted to public databases, with associated epidemiological metadata, from 10Tbp of raw sequencing data. The pipeline is robust to failure, auto restarting when one step fails. It estimates the amount of memory required. It performs multiple assemblies and several automated *in-silico* improvement steps that increase the contiguity of the resulting assembly. We assess the quality of the assemblies for low, neutral and high GC genomes. The pipeline is written in Perl and is freely available under the open source GNU GPL license.

## THEORY AND IMPLEMENTATION

An overview of the method is shown in Fig 1. For each genome, the *de novo* short-read assembler Velvet (Zerbino 2010) is used to generate multiple assemblies by varying the k-mer size between 66% and 90% of the read length using VelvetOptimiser (Gladman & Seemann 2008), as a well-chosen k-mer can substantially increase the quality of the resulting assembly (Zerbino 2010). From these assemblies, the assembly with the highest N50 is chosen. The N50 is the length L of the longest contig such that half of the nucleotides in the assembly lie in contigs of length at least L. When the pipeline was implemented the Velvet assembler was chosen because it proved to be robust to a wide range of data sets during testing and has a low computational overhead (Abbas et al. 2014) compared to SPAdes (Bankevich et al. 2012).

**Fig 1:**
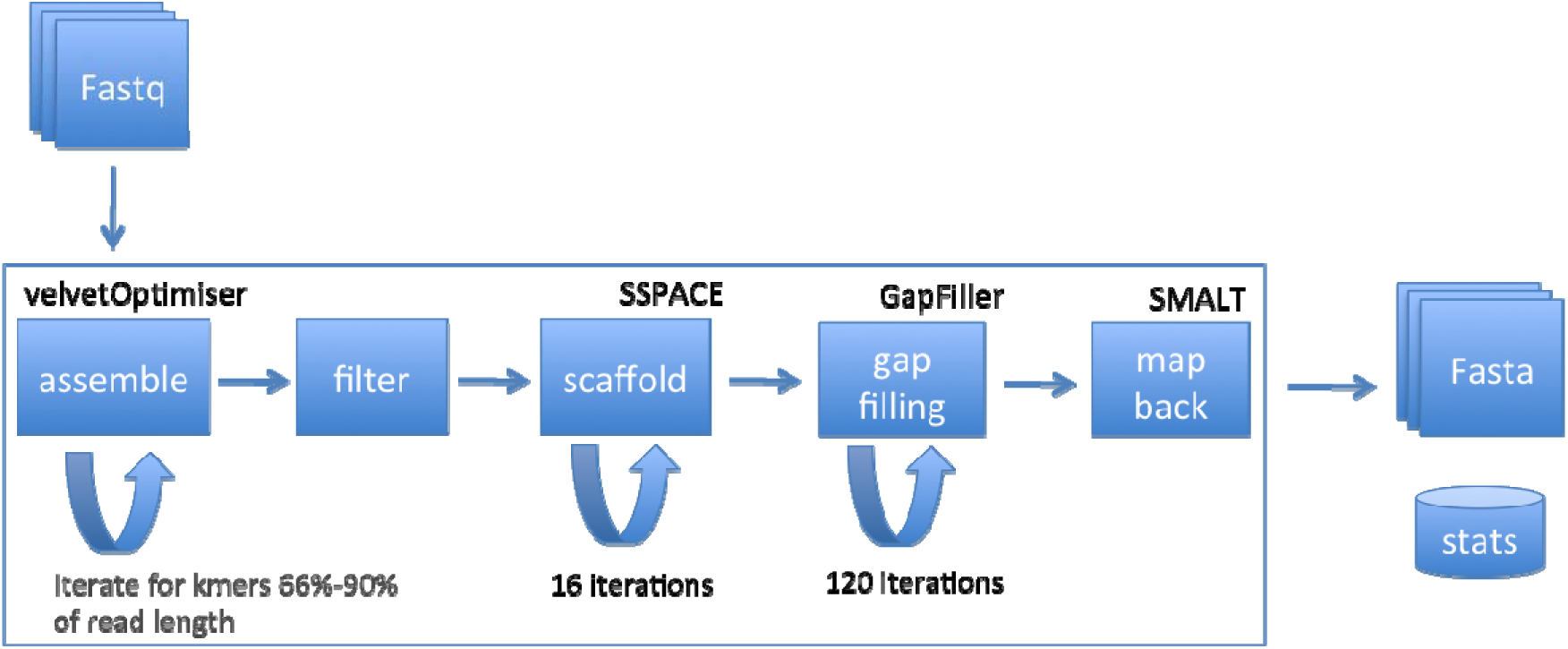
Overview of the method with major components noted.

A stand-alone assembly improvement step is run on the assembly to scaffold the contigs using SSPACE (Boetzer et al. 2011) and fill in sequence gaps using GapFiller (Boetzer & Pirovano 2012). First, to reduce the computational burden, reads that map (SMALT (Ponstingl & Ning 2015)) as proper pairs are excluded, since they have been successfully used in the assembly. A proper pair is a pair of reads from the same fragment of DNA which align to a single contig, in the correct orientation, within the expected insertion size range. The remaining reads, which are either unmapped or are mapped but have a mate that was unmapped or mapped to a different contig, are used for the improvement step. The contigs of the assembly are scaffolded by iteratively running SSPACE (Boetzer et al. 2011) (version 2.0), starting with pairs of contigs where there is the most read pair evidence. This reduces the likelihood of false joins by scaffolding contigs with the most linking information first. On the first iteration a minimum of 90 read pairs must link two contigs for them to be joined. This is then progressively reduced over 16 iterations down to 5 read pairs. These parameters were chosen after extensive testing on a range of organisms. Where two contigs are joined by read pairs, a gap consisting of an unknown number of bases (N) is generated. These gaps are targeted for closure by running 120 iterations of GapFiller (Boetzer & Pirovano 2012) (version 1.11), using a similar decreasing read evidence threshold beginning with a minimum depth of coverage of 90 reads, alternating between BWA (Li & Durbin 2009) and Bowtie (Langmead et al. 2009). Contigs are excluded from the assembly where they are shorter than the target fragment size (normally 300-500 bases). The contigs are then sorted by size and renamed in a standardised manner to include the raw sequencing data accession number. Finally, to assess the quality of the assembly and to produce a set of statistics, the reads are aligned again to the final assembly using SMALT. All the assemblies produced are created in a standardised manner and require no input from the user. The assemblies are then automatically annotated using PROKKA (Seemann 2014) with genus specific databases from RefSeq (Pruitt et al. 2012). The resulting annotated assemblies are in a format suitable for submission to EMBL/GenBank with post processing using GFF3toEMBL (Page et al. 2016). All the assemblies produced are created in a standardised manner and require no input from the user.

To assess the quality of the assemblies produced by the pipeline we used three microbial genomes with differing G+C content: *Bordetella pertussis* (67%), *Salmonella* Pullorum (52%) and *Staphylococcus aureus* (33%). This is a standard set of strains used for technology validation at the Wellcome Trust Sanger Institute (Quail et al. 2012). A closed complete capillary reference genome is available for each, with the *S. Pullorum* and *S. aureus* TW20 (Holden et al. 2010) data originating from the same isolate. Each were paired end sequenced on the Illumina MiSeq with a read length of 130bp, achieving a coverage of 28-43X. We compared the pipeline assemblies in each case to the capillary reference genomes using QUAST (Gurevich et al. 2013) and present the results in Table 1. Overall the assemblies contained at least 94% of the reference genome, so are good representations of the underlying genome. *S. Pullorum* was assembled into 22 contigs and *S. aureus* into 38 contigs. *B. pertussis* is known to contain many repetitive IS elements, explaining the higher level of fragmentation, which at 241 contigs is approximately equal to the number of IS elements (261 out of 3,816 genes in *B. pertussis* Tohama I annotated as IS elements). A pan genome was constructed using Roary (Page et al. 2015) for each organism, consisting of the predicted genes (Seemann 2014) from the reference and *de novo* assembly. The *de novo* assemblies contained 93-98% of the reference genes. This is in agreement with the percentage of the nucleotide bases matching between the *de novo* assembly and the reference, but does not account for misassemblies.

**Table 1:** Comparison of de novo assemblies derived from the pipeline against their corresponding complete reference genomes using QUAST.

| Organism | *B. pertussis* | *S. Pullorum* | *S. aureus* |
| --- | --- | --- | --- |
| Coverage | 40.16 | 28.11 | 43.86 |
| No. of contigs | 247 | 22 | 38 |
| Total length | 3,856,742 | 4,711,864 | 3,016,231 |
| Reference length | 4,086,189 | 4,895,678 | 3,075,806 |
| Genome fraction (%) | 94.32 | 95.74 | 98.00 |
| GC (%) | 67.81 | 52.15 | 32.64 |
| Reference GC (%) | 67.72 | 52.16 | 32.78 |
| N50 | 23,177 | 517,904 | 206,505 |
| No. of misassemblies | 6 | 10 | 4 |
| No. of mismatches per 100 kbp | 1.43 | 1.15 | 1.76 |
| No. of indels per 100 kbp | 0.6 | 1.92 | 0.17 |
| Genes | 3,610 | 4,460 | 2,814 |
| % reference genes found | 93.19 | 95.12 | 98.48 |

To assess the performance of the pipeline on a large scale we took a set of 18,080 published public assemblies and filtered them down to a set of 9,404 assemblies covering 73 bacterial species, summarised in Table 2. Only assemblies from isolates sequenced at the Wellcome Trust Sanger Institute on the Illumina HiSeq 2000/2500 or MiSeq platforms to high coverage (>50X) were considered. Contaminated samples were excluded after taxonomic classification of the raw reads with Kraken (Wood & Salzberg 2014). Fig. 2 gives the distribution of the number of contigs in each assembly. The mean is 89 contigs with peaks corresponding to different species, such as *Shigella* at 405 contigs. Before an isolate is sequenced, a reference genome is chosen based on the predicted species. We compared the size of the assembly to the size of the corresponding reference and present the distribution in Fig. 3. Ninety eight percent of all assemblies are within +/− 10% of the size of their corresponding reference genome. Some natural variation is to be expected within bacteria, for example the size of *Escherichia coli* genomes can vary by more than 20% (Blattner et al. 1997; Perna et al. 2001). Some may be larger because of plasmids or phage; others may have experienced gene loss and are smaller. However, most of the assemblies are at the expected size, allowing for useful comparisons to be made such as in (Wong et al. 2015; Makendi et al. 2016; Page et al. 2015).

**Table 2:** Summary of the isolates in the large public dataset.

| Species | No. Samples | Mean<br>Contigs | Mean Coverage |
| --- | --- | --- | --- |
| *Burkholderia pseudomallei* | 168 | 70 | 134 |
| *Campylobacter jejuni* | 379 | 24 | 121 |
| *Escherichia coli* | 178 | 167 | 145 |
| *Mycobacterium abscessus* | 157 | 37 | 120 |
| *Mycobacterium tuberculosis* | 1,441 | 122 | 150 |
| *Neisseria gonorrhoeae* | 234 | 75 | 205 |
| *Salmonella enterica* | 1,643 | 55 | 92 |
| *Salmonella Typhimurium* | 171 | 81 | 136 |
| *Shigella sonnei* | 299 | 405 | 118 |
| *Staphylococcus aureus* | 534 | 36 | 174 |
| *Staphylococcus haemolyticus* | 131 | 86 | 91 |
| *Streptococcus agalactiae* | 116 | 26 | 293 |
| *Streptococcus equi* | 159 | 81 | 374 |
| *Streptococcus pneumoniae* | 3,562 | 74 | 290 |
| Other | 232 | 80 | 136 |

**Fig 2:**
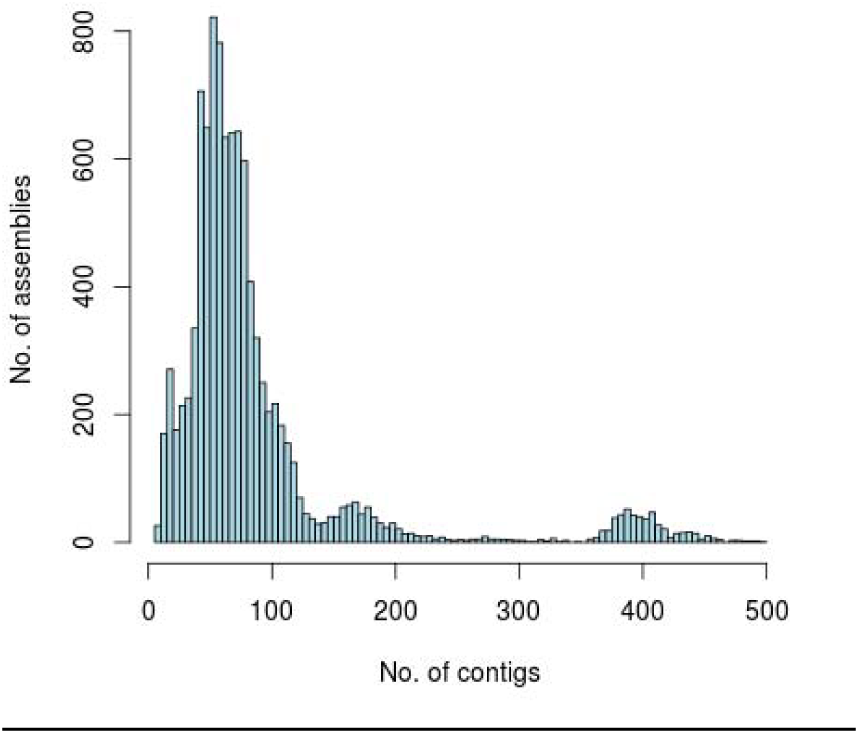
Distribution of the number of contigs in a set of 9,404 assemblies.

**Fig 3:**
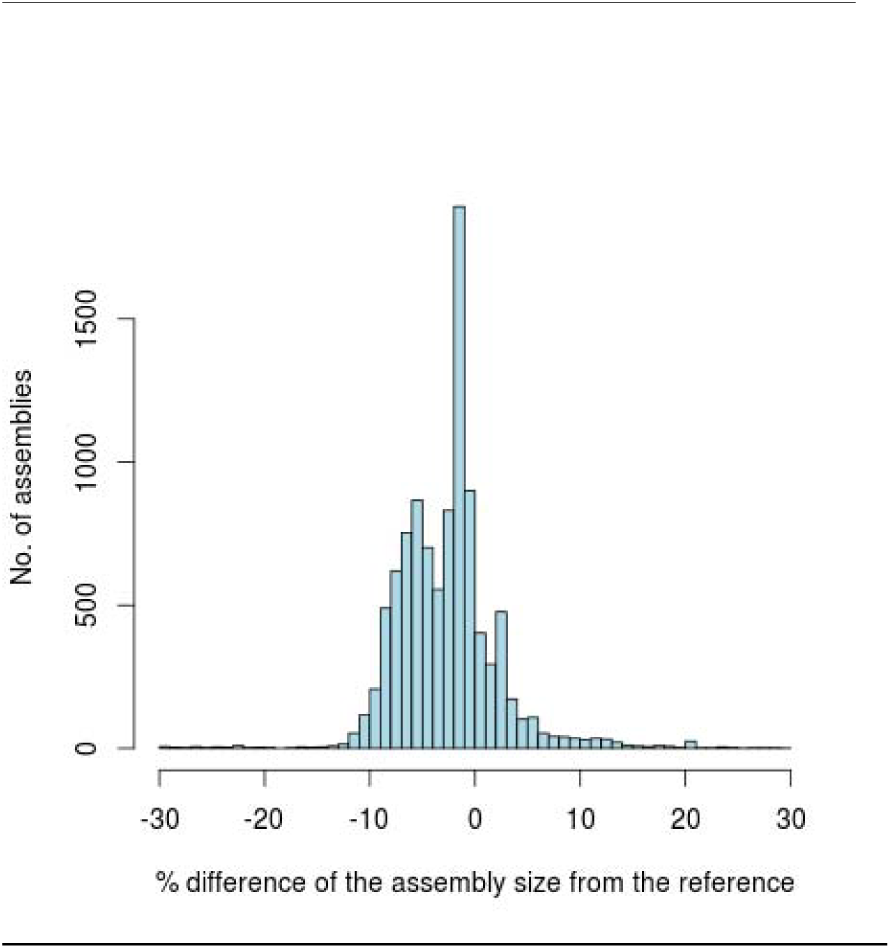
Fig 3: Distribution of the percentage difference between each assembly and the size of a closely related reference sequence.

Seven gene MLST schemes based on essential housekeeping genes exist for 6,971 of the assemblies (Maiden et al. 1998) from the set of 9,404 assemblies. These sequence typing methods are widely used by reference labs for genomic epidemiology, predating whole genome sequencing technologies. If all of the MLST genes are present in the assemblies then it allows for the assemblies to be used as a replacement for traditional PCR based methods. The MLST scheme for *Mycobacterium abscessus* is poorly populated, containing very few alleles and we could only assign an allele in 30% of cases, so has been excluded from this analysis, leaving 6,814 assemblies. Only genes with at least 95% length and identity to a known MLST allele are counted as a match. We found that in 6,789 (99.6%) assemblies we could identify all of the MLST genes using MLST-check (Page 2016), a method which performs a nucleotide blast (Camacho et al. 2009) of all the MLST alleles against each assembly, with the latest databases downloaded from pubMLST (Jolley & Maiden 2010). Sixteen assemblies were missing 1 MLST gene (0.23%). One sample (0.013%) was only partially assembled but on closer investigation it had unusually high coverage (445X), which appears to have lead to a poor choice of *k*-mer. Of the remaining 8 assemblies, where the sequence type could not be inferred from the assembly, all contained contamination were identified as different species when analysed with Kraken (Wood & Salzberg 2014).

## CONCLUSION

Generating annotated genomes from whole genome sequencing data is a complex and laborious process that enables the true diversity within a species to be unveiled. We developed a high throughput pipeline that has been used to generate 30% of all bacterial assemblies in GenBank. The resulting genomes encompass more than 94% of the predicted genes and nucleotides, and have MLST genes available in 99.6% of assembled samples over a range of organisms with different GC content. We demonstrate that it has been successfully scaled up to tens of thousands of samples, providing annotated *de novo* assemblies suitable for submission to EMBL/GenBank without the need for manual intervention.

## ACKNOWLEDGEMENTS

This work was supported by the Wellcome Trust (grant WT 098051).

## ABBREVIATIONS

MLST: Multilocus sequence typing

